# Autophagy contributes to homeostasis in esophageal epithelium where high autophagic vesicle content marks basal cells with limited proliferation and enhanced self-renewal potential

**DOI:** 10.1101/2023.09.20.558614

**Authors:** Alena Klochkova, Adam L. Karami, Annie D. Fuller, Louis R. Parham, Surali R. Panchani, Shruthi Natarajan, Jazmyne L. Jackson, Anbin Mu, Yinfei Tan, Kathy Q. Cai, Andres J. Klein-Szanto, Amanda B. Muir, Marie-Pier Tétreault, Kathryn E. Hamilton, Kelly A. Whelan

## Abstract

**Background & Aims:** Autophagy has been demonstrated to play roles in esophageal pathologies both benign and malignant. Here, we aim to define the role of autophagy in esophageal epithelium under homeostatic conditions.

**Methods:** We generated tamoxifen-inducible, squamous epithelial-specific *Atg7* (autophagy related 7) conditional knockout mice to evaluate effects on esophageal homeostasis and response to the carcinogen 4-nitroquinoline 1-oxide (4NQO) using histological and biochemical analyses. We FACS sorted esophageal basal cells based upon fluorescence of the autophagic vesicle (AV)-identifying dye Cyto-ID, then subjected these cells to transmission electron microscopy, image flow cytometry, 3D organoid assays, RNA-Sequencing (RNA-Seq), and cell cycle analysis. 3D organoids were subjected to passaging, single cell (sc) RNA-Seq, cell cycle analysis, and immunostaining.

**Results:** Genetic autophagy inhibition in squamous epithelium resulted in increased proliferation of esophageal basal cells. Esophageal basal cells with high AV level (Cyto-ID^High^) displayed limited organoid formation capability upon initial plating but passaged more efficiently than their counterparts with low AV level (Cyto-ID^Low^). RNA-Seq suggested increased autophagy in Cyto- ID^High^ esophageal basal cells along with decreased cell cycle progression, the latter of which was confirmed by cell cycle analysis. scRNA-Seq of 3D organoids generated by Cyto-ID^Low^ and Cyto- ID^High^ cells identified expansion of 3 cell populations, enrichment of G2/M-associated genes, and aberrant localization of cell cycle-associated genes beyond basal cell populations in the Cyto- ID^High^ group. Ki67 expression was also increased in organoids generated by Cyto-ID^High^ cells, including in cells beyond the basal cell layer. Squamous epithelial-specific autophagy inhibition induced significant weight loss in mice treated with 4NQO that further displayed perturbed epithelial tissue architecture.

**Conclusions:** High AV level identifies esophageal epithelium with limited proliferation and enhanced self-renewal capacity that contributes to maintenance of the esophageal proliferation- differentiation gradient *in vivo*.

## Introduction

Esophageal epithelium is a stratified squamous tissue. Under homeostatic conditions, basal keratinocytes proliferate then undergo passive migration toward the luminal surface before desquamating into the lumen. Creating a proliferation-differentiation gradient at an exquisite level, the described dynamic process further permits epithelial renewal which occurs over a period of ∼2 weeks^1^. Maintenance of the esophageal epithelial proliferation-differentiation gradient is critical as esophageal epithelium is the first line barrier to prevent penetration of digestive contents. Disruption of this gradient is associated with widely prevalent esophageal pathologies, including esophagitis and cancer, where basal cell hyperplasia (BCH) is a common epithelial remodeling event. Molecular markers defining basal keratinocytes include cell surface molecules integrin β1 (CD29), integrin α6 (CD49f), transferrin receptor (CD71), and neurotrophin receptor p75NTR, cytokeratins KRT5 and KRT14 and transcription factors p63 and SOX2^2–7^. Single cell RNA-Sequencing (scRNA-Seq) studies in human and murine esophageal epithelium have revealed marked cellular heterogeneity within esophageal epithelium and particularly in basal cells^8–15^. Several studies have further demonstrated functional heterogeneity among murine esophageal basal cells linked to expression of specific markers, including CD34, KRT15, and CD73^16–18^. However, studies utilizing lineage tracing in mice coupled with mathematical modeling have proposed a single-progenitor model in esophageal epithelium wherein all basal cells have equal capacity to proliferate and differentiate ^1, 19^.

Autophagy (macroautophagy) is a catabolic process that mediates lysosomal degradation of intracellular components via double-membraned autophagic vesicles (AVs). The role of autophagy in esophageal biology is an emerging area of interest. Numerous studies have evaluated autophagy in relation to esophageal cancer, both esophageal squamous cell carcinoma (ESCC) and esophageal adenocarcinoma (EAC)^20^. Overall, this body of literature supports various roles for autophagy in esophageal malignancy, including supporting cancer stem cell (CSC) biology. We have shown that autophagy promotes expansion of ESCC CSCs in response to TGF-β^21^. The core autophagy protein autophagy related (ATG)7 has also been demonstrated to support ESCC CSC maintenance by promoting β-catenin stabilization^22^. In addition to roles for autophagy in cancer, our own findings demonstrate that autophagy is a critical cytoprotective mechanism in esophageal epithelium upon exposure to stressors relevant to eosinophilic esophagitis^23^ and gastroesophageal reflux disease/Barrett’s esophagus^24^, highlighting the potential for broad functional significance of this evolutionarily conserved pathway in esophageal biology.

Here, we explore the role of autophagy in esophageal epithelium under homeostatic conditions. We demonstrate that genetic depletion of autophagy disrupts the homeostatic proliferation-differentiation gradient of esophageal epithelium *in vivo*. We show that esophageal basal cells with high AV level are less proliferative than their counterparts with low AV level. Conversely, esophageal basal cells with high AV level exhibit increased self-renewal in 3D organoid assays. scRNA-Seq coupled with immunostaining for Ki67 further demonstrate aberrant proliferation in 3D organoids generated by basal cells with high AV level. Finally, we report that genetic depletion of autophagy in squamous epithelial cells is deleterious in mice exposed to the oral-esophageal carcinogen 4-nitroquinoline 1-oxide (4NQO).

## Results

### Conditional ATG7 depletion disrupts the homeostatic proliferation-differentiation gradient of murine esophageal epithelium

To investigate the functional role of autophagy in esophageal homeostasis, we generated *K5CreERT2^wt/mut^; Atg7^loxp/loxp^ mice* with tamoxifen (TAM)-inducible conditional depletion of the essential autophagy gene *Atg7* in K5-expressing cells (i.e. basal cells) of squamous epithelium. K5CreERT2^wt/mut^; Atg7^loxp/loxp^ mice were treated with TAM by oral gavage at the start of the experiment and then again 2 weeks later (**Figure 1A**), resulting in decreased expression in ATG7 at the level of RNA and protein in esophageal mucosa 6 weeks following the initial TAM treatment (∼4-6 turnover cycles) (**Figure 1B, C**). ATG7 depletion promoted increased thickness of the combined epithelium and overlying keratin layer (**Figure 1D-E**), indicating perturbation of the homeostatic proliferation/differentiation gradient. Ki67 labeling further confirmed increased number of proliferative cells in the basal cell layer of esophageal epithelium (**Fig. 1F-G**). These findings suggest that autophagy contributes to maintenance of the proliferation-differentiation gradient of esophageal epithelium under homeostatic conditions.

**Figure 1.**
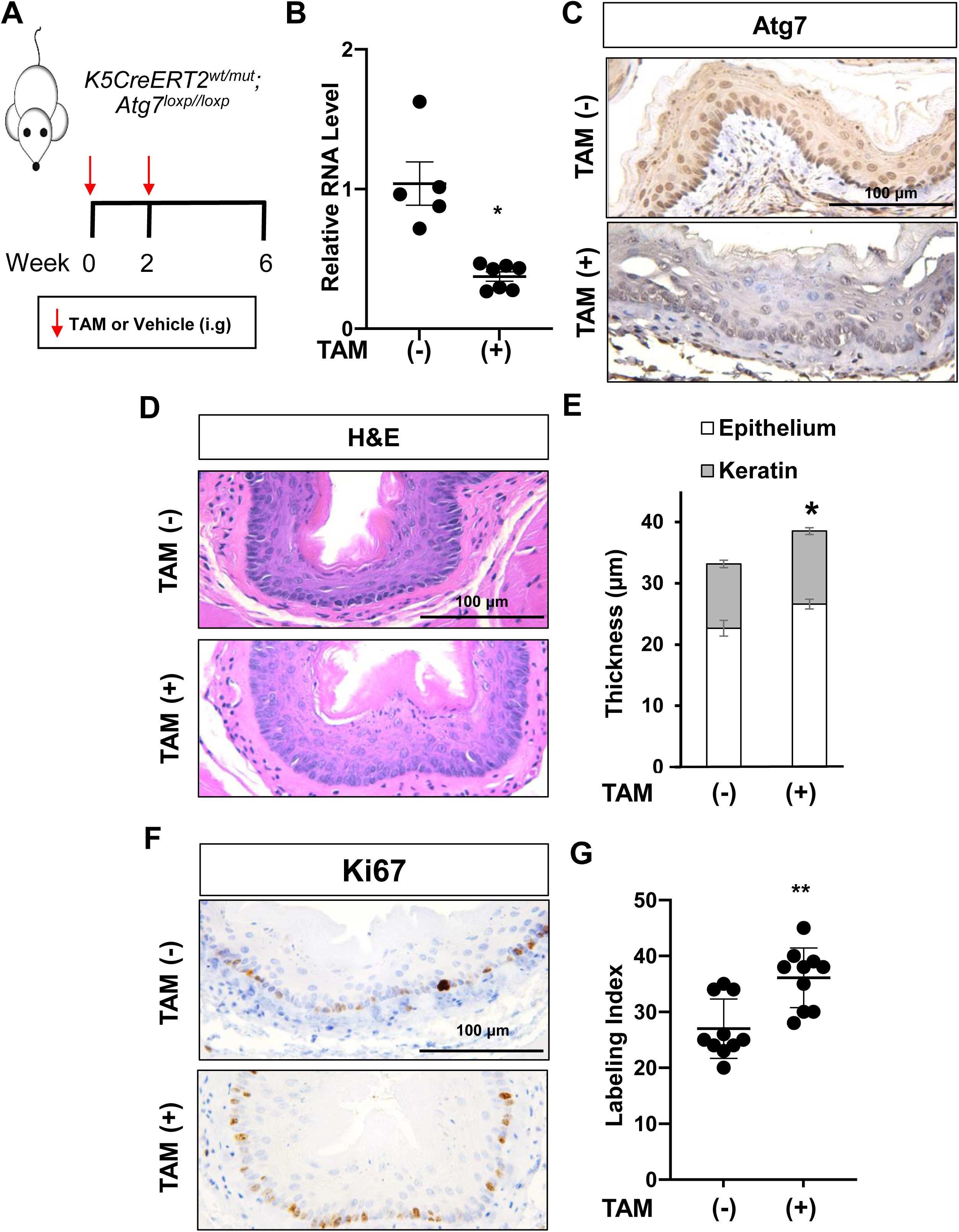
Effects of genetic Atg7 depletion in esophageal epithelium. (A) Schematic overview of experimental design. K5Cre^ERT2+/-^; Atg7^loxp/loxp^ mice were gavaged with vehicle or tamoxifen (TAM) at indicated time points. Esophagi were harvested 6 weeks after initial dose of TAM. **(B)** Dot plot showing relative RNA expression of *Atg7* in epithelium-enriched portion of esophagus. Each dot represents an individual mouse, bars represent Mean±SEM (*p<0.05 by t-test). **(C)** Representative ATG7 staining of esophageal epithelium. Scale bar, 100 μm. **(D)** Representative H&E staining of esophageal epithelium. Scale bar, 100 μm. **(E)** Thickness of epithelium measured for H&E sections of esophagi cut in the traverse plane. Bar diagram shows Mean±SEM for epithelium and overlying keratin layer (n=10; *p<0.05 by t-test). **(F)** Representative ATG7 staining of esophageal epithelium. Scale bar, 100 μm. **(G)** Dot plot showing Ki67+ labeling index in esophageal epithelium. Each dot represents mean index for an individual mouse, bars represent Mean±SEM (n=10; **p<0.005 by t-test).

### Esophageal basal cells with high AV level display diminished proliferation coupled with enhanced self-renewal potential

We have previously demonstrated evidence of AVs in esophageal basal cells of both humans and mice^23, 24^. These findings in conjunction with increased proliferation in the basal cell layer of esophageal epithelium from ATG7-depleted mice led us to investigate the relationship between AV level and esophageal basal cell biology. To that end, we used FACS to isolate esophageal basal cells with high or low AV level based upon the expression of the basal marker Integrin β1 and the fluorescence of the AV-identifying dye Cyto-ID (**Figure 2A**). After confirming that esophageal basal cells with high Cyto-ID fluorescence (Cyto-ID^High^ cells) displayed increases in both discrete Cyto-ID puncta and AVs as compared to their counterparts with low levels of Cyto-ID fluorescence (Cyto-ID^Low^ cells) (**Figure 2B, C**), we subjected these sorted populations to 3D organoid assays. We found that Cyto-ID^Low^ cells more readily formed organoids compared to Cyto- ID^High^ cells (**Figure 2D, E**). We further observed marked differences in morphology when comparing organoids generated by Cyto-ID^Low^ and Cyto-ID^High^ cells (**Figure 2F-H**). The majority (80.3%) of organoids generated by Cyto-ID^Low^ basal cells displayed 1-4 layers of basal cells at the smooth outer periphery, several inner layers of cells displaying flattened morphology, and a central keratinized core, consistent with terminal squamous differentiation (**Figure 2F-H**). By contrast this ‘regular’ phenotype was detected in 45.4% of organoids generated by Cyto-ID^High^ basal cells. Instead, organoids derived from Cyto-ID^High^ basal cells were enriched for ‘irregular’ phenotypes that included increased basal cell content and the presence of multiple basal-cell rich areas protruding at the structure’s outer edges (present in 31.7% organoids) **(Figure 2F-H**), the latter of which are reminiscent of “buds” that have been associated with stemness in intestinal enteroid culture^25^. 96.9% of organoids generated by Cyto-ID^High^ cells displayed evidence of keratinization, suggesting that these cells are capable of terminal differentiation. In order to evaluate self-renewal capacity, we continued to passage organoids derived from Cyto-ID^Low^ and Cyto-ID^High^ esophageal basal cells. Although Cyto-ID^Low^ cells formed organoids more readily upon initial plating, their ability to form consistently declined over passaging with a total loss of detectable organoids following passage 2 (**Figure 2D, E**). By contrast, Cyto-ID^High^ basal cells initially displayed limited ability to generate organoids but maintained passaging out to passage 4 (**Figure 2D, E**).

**Figure 2.**
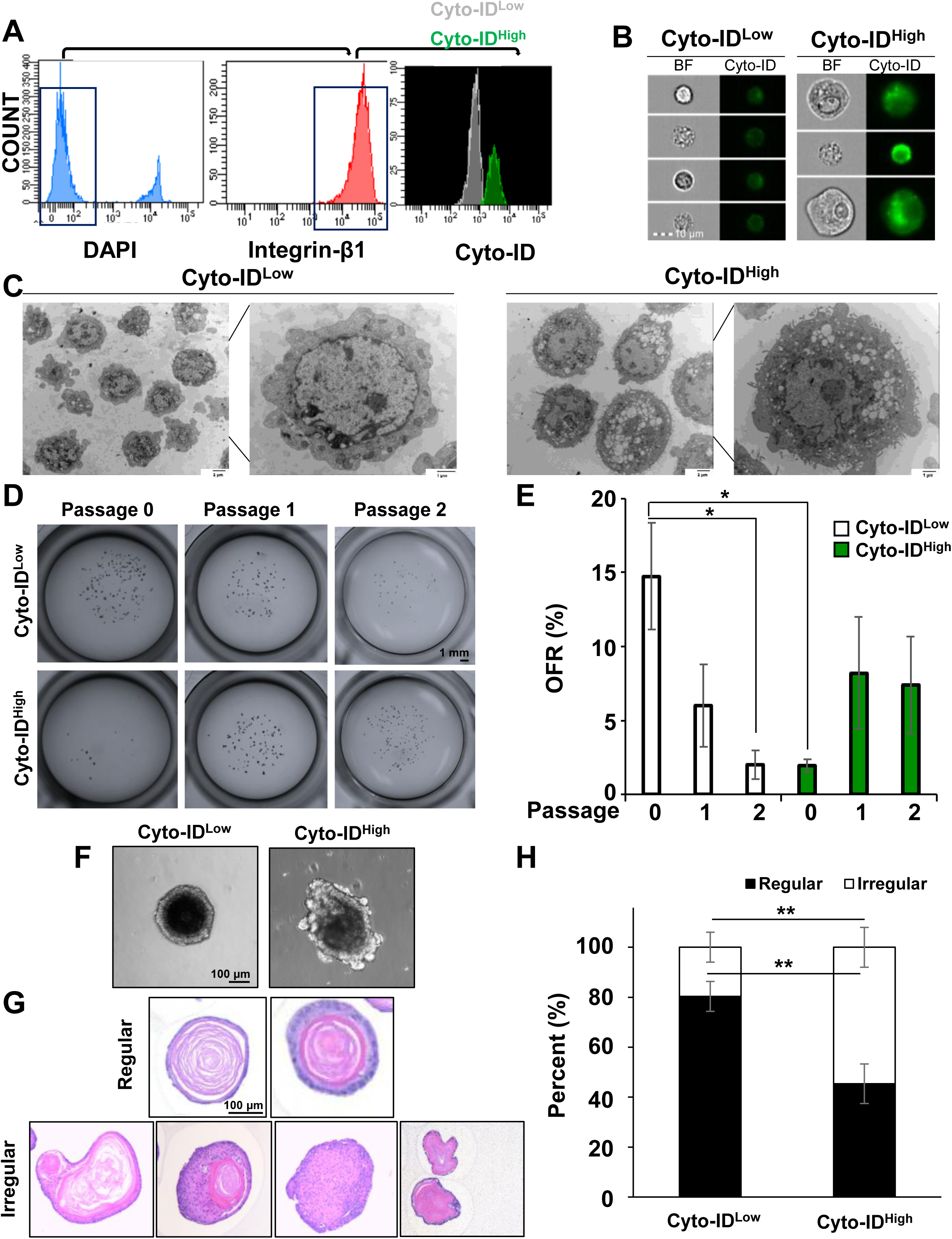
Characterization of Cyto-ID^Low^ and Cyto-ID^High^ basal cells from murine esophageal epithelium. (A) FACS gating strategy used to isolate live (DAPI^negative^) Cyto-ID^Low^ and Cyto-ID^High^ basal cells (Integrinβ1^positive^). **(B)** Representative images of Cyto-ID^Low^ and Cyto-ID^High^ cells acquired using imaging flow cytometry. Bar 10 μm (n=3). **(C)** Representative TEM images of sorted Cyto-ID^Low^ and Cyto-ID^High^ cells. (**D-H**) Cyto-ID^Low^ and Cyto-ID^High^ basal cells were sorted from mouse esophageal epithelium then cultured in 3D organoid assays for 13-15 days. **(D)** Representative images of organoids formed either directly after sorting (passage 0) or following passaging. For passaging, organoids were dissociated after 13-15 days in culture and resulting single cell suspensions were plated in 3D organoid assays. **(E)** Organoid formation rate (OFR; number of organoids formed/number of cells plated x 100) was determined for Cyto-ID^Low^ and Cyto-ID^High^ basal cells at indicated passages. Data is presented as Mean±SEM (n=4, *p<0.05 by t-test). **(F)** Representative bright field images of Cyto-ID^Low^ and Cyto-ID^High^ organoids (passage 0). **(G, H)** H&E-stained organoids were evaluated for morphology. Representative images of regular and irregular organoids are shown in **(G)** and bar diagram in **(H)** shows distribution of regular and irregular organoids generated by organoids Cyto-ID^Low^ and Cyto-ID^High^ cells (n=9, **p<0.01 by t- test).

We continued to explore the molecular features of Cyto-ID^High^ and Cyto-ID^Low^ esophageal basal populations by performing RNA-Seq on these sorted populations. Principal component analysis revealed sample separation based on Cyto-ID level, but not by sex (**Figure 3A**). We identified 7,102 differentially expressed genes (DEGs) when comparing Cyto-ID^Low^ and Cyto-ID^High^ cells (**Figure 3B**; **Supplementary File S1**). IPA further predicted alterations in 110 pathways (**Supplementary File S1**) with Autophagy being among the top 10 upregulated pathways (**Figure 3C**). Cell Cycle Control of Chromosomal Replication and Kinetochore Metaphase Signaling Pathway were the top 2 pathways predicted to be downregulated in Cyto-ID^High^ cells (**Figure 3C**). As limited cell proliferation may contribute to the diminished organoid formation capacity observed in Cyto-ID^High^ cells (**Figure 2D, E**), we next used flow cytometry analysis for PI to assess cell cycle in Cyto-ID^High^ and Cyto-ID^Low^ esophageal basal cells. In both populations, cells in the G0/G1 phase of the cell cycle were predominant (**Figure 4A, B**), which is in agreement with published literature in primary esophageal keratinocytes^26^. Cyto-ID^High^ cells, however, displayed an increase in the G0/G1 fraction coupled with depletion of cells in the S and G2/M phases of the cell cycle (**Figure 4A, B**).

**Figure 3.**
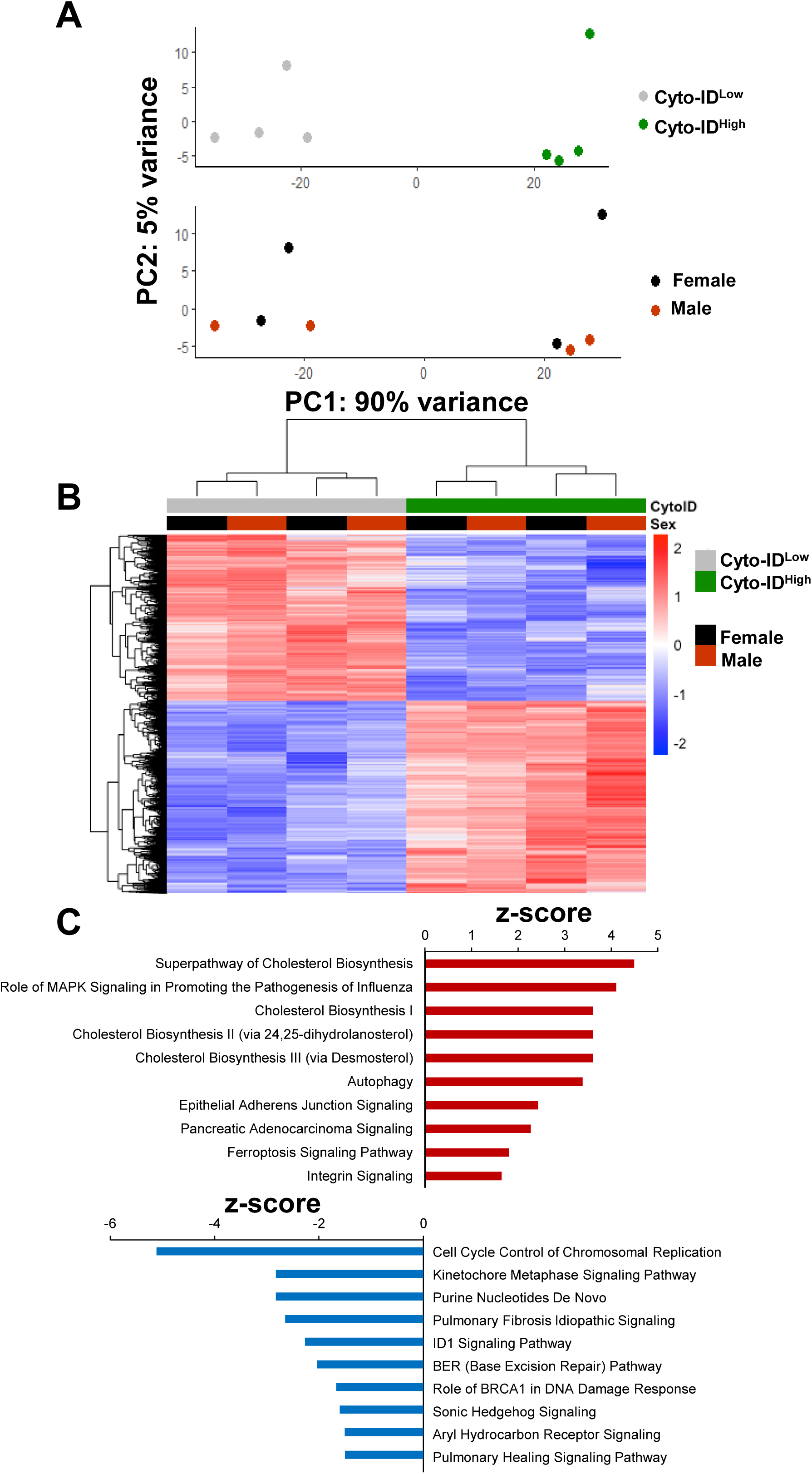
Transcriptomic analysis of Cyto-ID^Low^ and Cyto-ID^High^ basal cells. Cyto-ID^Low^ and Cyto-ID^High^ basal cells from 4 independent experiments were sorted from mouse esophageal epithelium then subjected to RNA-Seq. **(A)** Principal Component Analysis showing clustering of samples according to Cyto-ID level (top) and sex (bottom). **(B)** Heatmap showing relative expression pattern in Cyto-ID^High^ vs Cyto-ID^Low^ cells (p≤0.01). **(C)** Top 10 significantly dysregulated canonical pathways based on Ingenuity Pathway Analysis (IPA) z-score. Red indicates enrichment and blue indicates inhibition.

**Figure 4.**
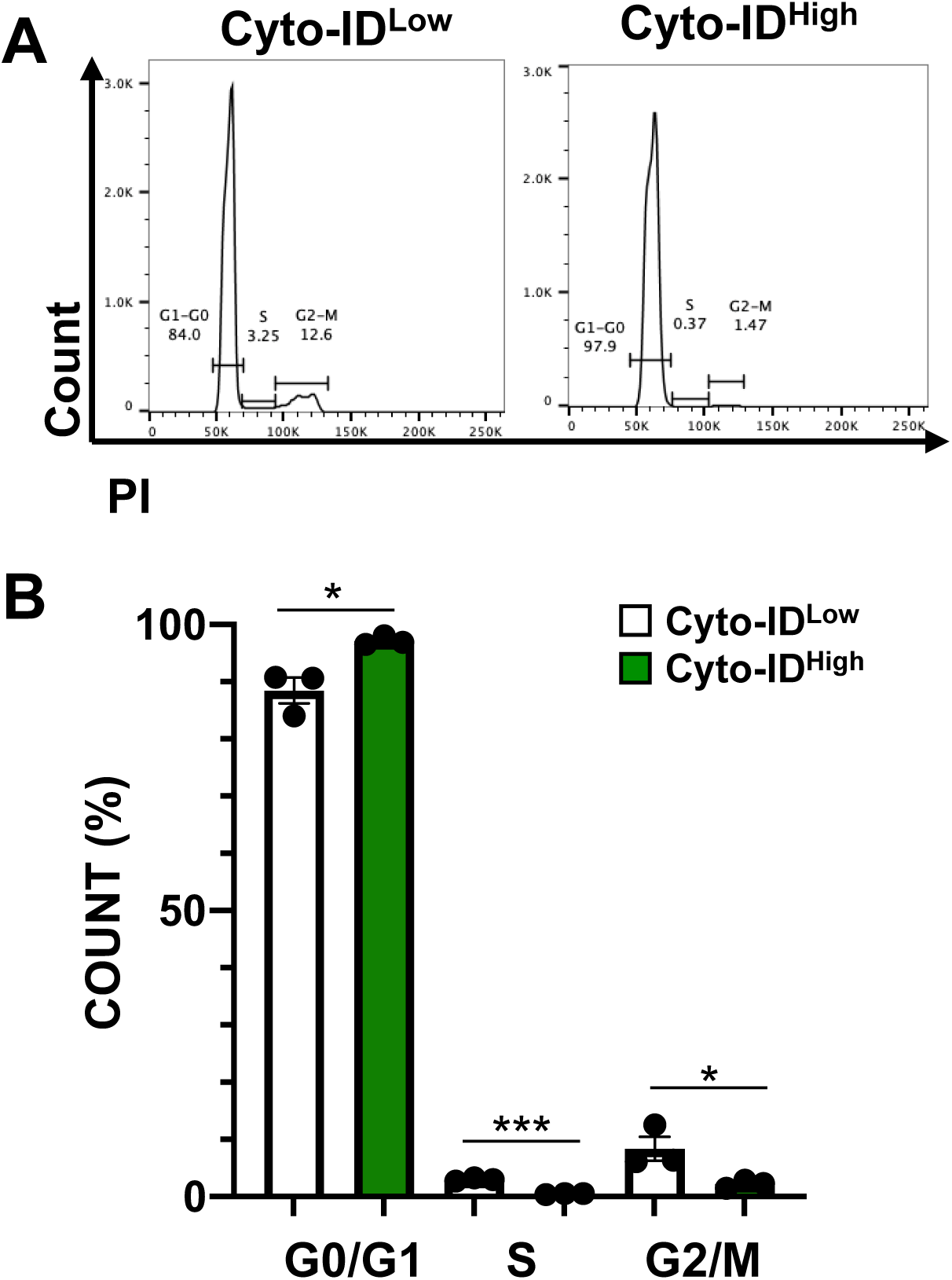
Cell cycle evaluation in Cyto-ID^Low^ and Cyto-ID^High^ esophageal basal cells. Cyto- ID^Low^ and Cyto-ID^High^ basal cells were sorted from mouse esophageal epithelium then subjected to DNA content analysis using propidium iodide (PI) flow cytometry. **(A)** Representative histogram of PI staining. **(B)** Bar diagram shows average distribution of cells in each phase of the cell cycle. Data is presented as Mean±SEM (n=3; *p<0.05; ***p≤0.0001 by t-test).

### 3D organoids generated by esophageal basal cells with high AV level feature increased proliferation that extends beyond the basal cell layer

As Cyto-ID^Low^ and Cyto-ID^High^ esophageal basal cells displayed marked differences in morphology and passaging capability, we next evaluated the cellular and molecular heterogeneity of these cells using scRNA-Seq (**Figure 5A; Supplementary Figure S1**). We identified 13 cell populations in organoids generated by Cyto-ID^Low^ and Cyto-ID^High^ esophageal basal cells (**Figure 5B, C**). We evaluated expression of the basal cell marker *Krt5* (encoding Keratin 5) the superficial cell marker *Krt13* (encoding Keratin 13) in our dataset in order to establish the basal-superficial cell axis (**Figure 5D-F**). In doing so, we identified 6 basal, 1 suprabasal, and 6 superficial populations. We then evaluated the representation of each cell type in organoids generated by Cyto-ID^Low^ and Cyto-ID^High^ esophageal basal cells, finding a significant increase in the abundance of 3 populations in the Cyto-ID^High^ group: basal 2, basal 3, and superficial 1 (**Figure 6A, B**). Basal 3 was of particular interest as pathway analysis suggested activation of the cell cycle in this cell population (**Figure 7**). We continued to map expression of cell cycle-associated genes onto our dataset, revealing enrichment of G2/M genes in population basal 3 and S phase genes in population basal 2 (**Figure 8A**). scRNA-Seq further identified an increase in G2/M genes in organoids generated by Cyto-ID^High^ esophageal basal cells as compared to those generated by their Cyto-ID^Low^ counterparts (**Figure 8B**). Flow cytometry for PI in organoids derived from Cyto- ID^Low^ and Cyto-ID^High^ esophageal basal cells validated increased G2/M cell in the Cyto-ID^High^ group (**Figure 8C, D**). Ki67 staining was also increased in organoids derived from Cyto-ID^High^ esophageal basal cells (**Figure 8E, F**). Furthermore, both expression of G2/M genes and staining for Ki67 was located beyond the basal cell compartment in organoids derived from Cyto-ID^High^ esophageal basal cells (**Figure 8A, E**). We also found that Cyto-ID fluorescence in organoids generated by Cyto-IDLow and Cyto-IDHigh cells displayed comparable levels of Cyto-ID fluorescence (**Figure 8G, H**). These data indicate that although Cyto-ID^High^ basal cells within esophageal epithelium exhibit limited proliferation, the organoids generated by these Cyto-ID^High^ cells feature increased proliferative capability, including in cells that are not part of the basal cell layer, and fail to maintain elevated AV level.

**Figure 5.**
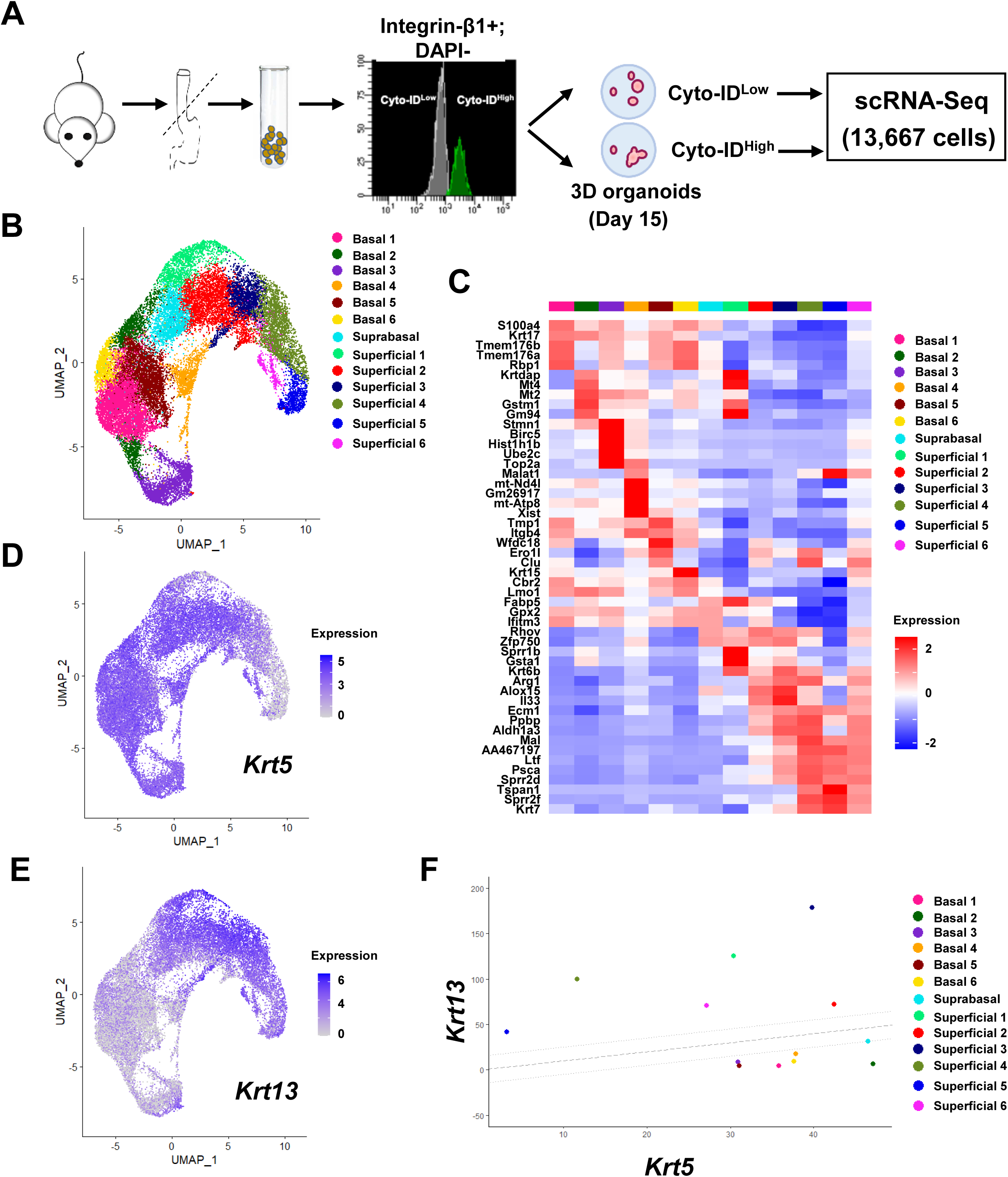
Identification of cell populations in 3D organoids generated by Cyto-ID^Low^ and Cyto-ID^High^ murine esophageal basal cells. (A) Schematic overview of experimental design. **(B)** Seurat’s Uniform Manifold Approximation and Projection (UMAP) was used to identify 13 cell populations within the scRNA-Seq dataset. (**C)** Expression z-scores for the top 5 most upregulated genes in each cluster. Red indicates enrichment while blue indicates inhibition. **(D,E)** Log1p normalized expression of the basal marker *Krt5* (D) and superficial marker *Krt13* (E), across the dataset is shown. Purple indicates enrichment. **(F)** Normalized log10 expression of *Krt5* vs normalized log10 expression of *Krt13* was plotted for all populations identified in the scRNA-Seq dataset. Populations falling within the strip defined by lines with slope of 1 passing through points (*0, -15*) and (*0, 15*) were define as suprabasal. Populations falling below this strip were defined as basal and those above this strip were identified as superficial.

**Figure 6.**
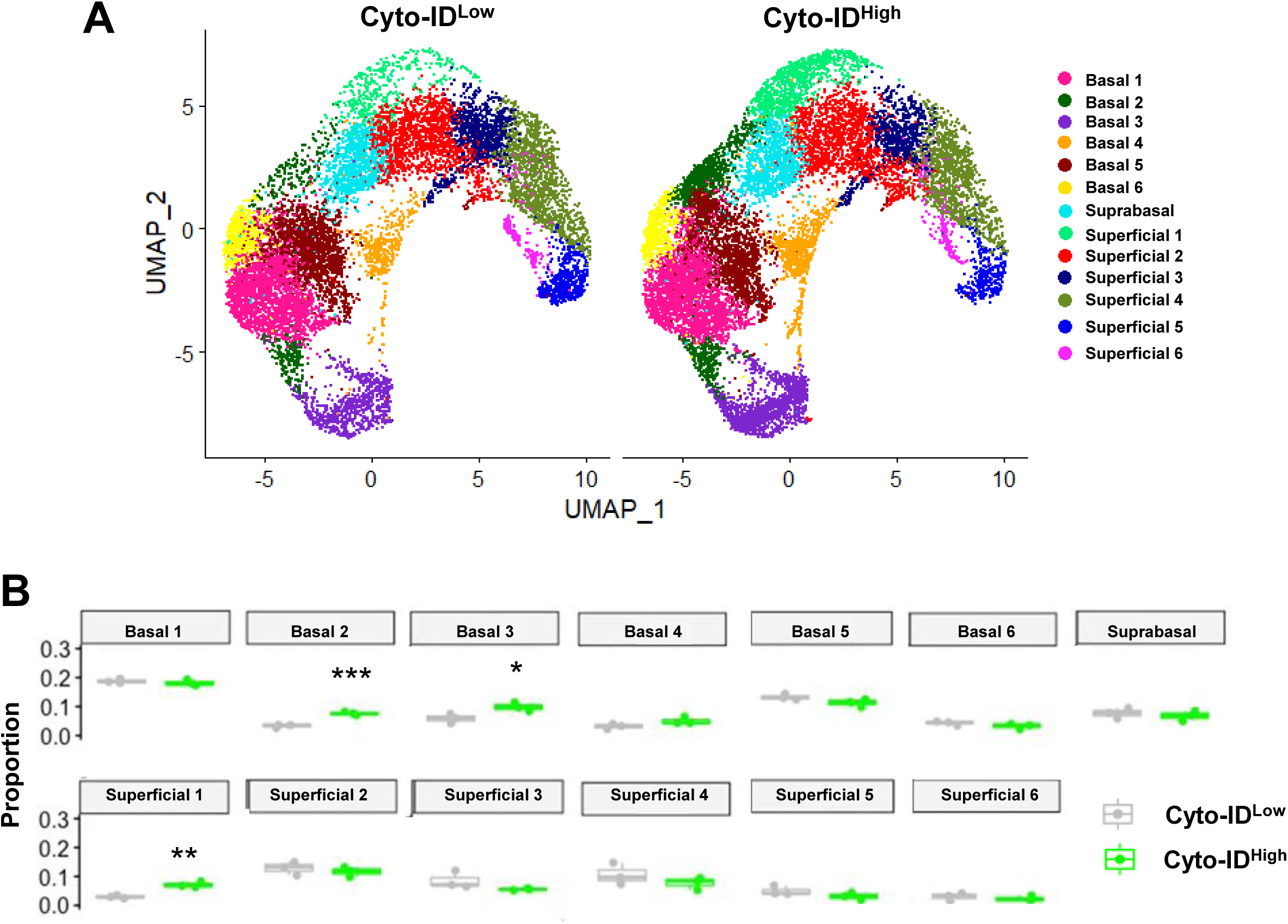
Representation of 13 cell populations in organoids generated by Cyto-ID^Low^ and Cyto-ID^High^ murine esophageal basal cells. scRNA-Seq was performed in organoids generated by Cyto-ID^Low^ and Cyto-ID^High^ basal cells from mouse esophageal epithelium. **(A)** Seurat’s Uniform Manifold Approximation and Projection (UMAP) was used to assess distinct cell populations in organoids generated by Cyto-ID^Low^ and Cyto-ID^High^ murine esophageal basal cells. **(B)** Box plots showing proportion of each cell as a fraction of all cells for each cluster in Cyto-ID^High^ organoids compared to Cyto-ID^Low^, where each dot represents one biological repeat (*p<0.05*; p≤0.01**; ***p<0.001 by t-test).

**Figure 7.**
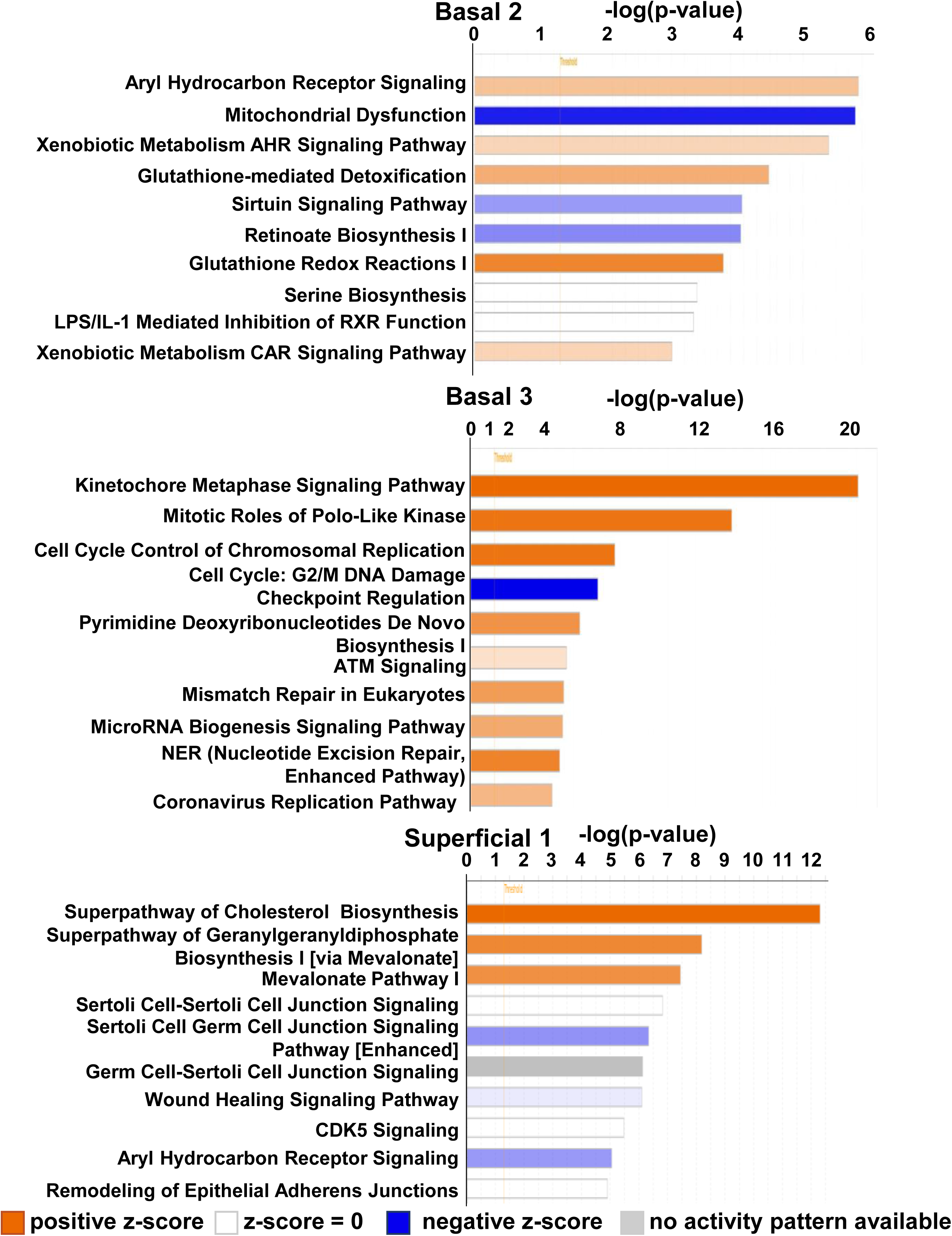
Ingenuity Pathway Analysis of Basal 2, 3 and Superficial 1 cell populations. scRNA-Seq was performed in organoids generated by Cyto-ID^Low^ and Cyto-ID^High^ basal cells from mouse esophageal epithelium. Top 10 cellular processes predicted to be significantly altered based on Ingenuity Pathway Analysis (IPA) z-score in indicated cell populations (threshold=1.3). Orange indicates enrichment and blue indicates inhibition.

**Figure 8.**
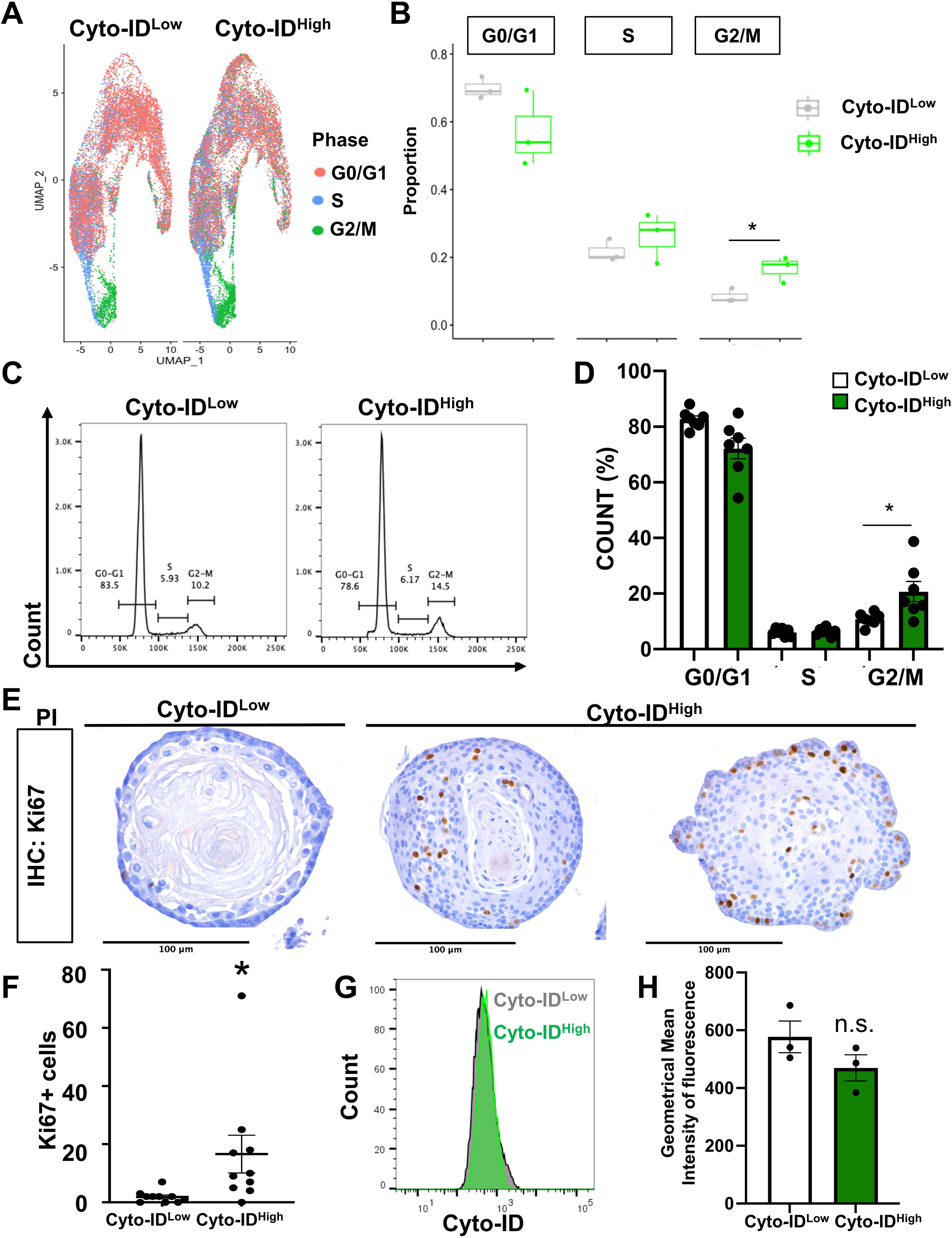
Assessment of proliferation in organoids generated by Cyto-ID^Low^ and Cyto-ID^High^ esophageal basal cells. Cyto-ID^Low^ and Cyto-ID^High^ basal cells were sorted from mouse esophageal epithelium and cultured in 3D organoid assays for 15 days. **(A, B)** scRNA-Seq analysis generated Seurat’s Uniform Manifold Approximation and Projection (UMAP) onto which expression of genes associated with each cell cycle phase was projected in **(A**). In **(B),** box plots show predicted portion of cells from Cyto-ID^Low^ and Cyto-ID^High^ organoids in each cell-cycle phase. Each dot represents one biological repeat (*p<0.05 by t-test). **(C-F)** Cyto-ID^Low^ and Cyto-ID^High^ basal cells were cultured in 3D organoid assays for 15 days. Organoids were dissociated then subjected to DNA content analysis via propidium iodide (PI) flow cytometry with representative histogram shown in **(C)** and bar diagram of average distribution of cells in each phase of the cell cycle shown in **(D)**. Data in **(D)** is presented as Mean±SEM (n=7; *p<0.05 by t-test). Organoids were stained for Ki67 by immunohistochemistry in **(E)** with number of Ki67-positive cells per organoid shown in **(F)**. In **(F)**, each dot represents one organoid and bars represent Mean±SEM (*p<0.05). **(G, H)** Organoids were dissociated and stained for Cyto-ID with representative flow plots shown in **(G)** and quantification of Cyto-ID fluorescence shown in **(H)** with each dot representing 1 biological repeat (n.s. by t-test)).

### ATG7 in squamous epithelial cells is indispensable during response to 4NQO challenge

As we identified diminished proliferation and increased self-renewal potential in esophageal basal cells with high AV level, we finally sought to determine the effects of autophagy depletion in esophageal epithelium responding to a stimulus that drives proliferation. After confirming that TAM treatment reduced the percentage of Cyto-ID^High^ cells in esophageal epithelium of *K5CreERT2^wt/mut^; Atg7^loxp/loxp^* mice (**Figure 9A**), we challenged these mice with 4NQO, a potent oral-esophageal carcinogen that induces proliferation in esophageal epithelium^27–29^. We planned to treat mice with 4NQO for 16 weeks to assess effects of genetic autophagy inhibition in the context of esophageal dysplasia. However, after only 14 days of 4NQO, complications arose in TAM-treated *K5CreERT2^wt/mut^; Atg7^loxp/loxp^* mice, where ATG7 mRNA depletion was apparent (**Figure 9B, C**). Of the 5 TAM-treated *K5CreERT2^wt/mut^; Atg7^loxp/loxp^*mice that were administered 4NQO, 2 died abruptly while the remaining 3 mice exhibited significant decline in body weight (**Figure 9D**). Histological evaluation further revealed BCH coupled with cytoplasmic vacuolization of basal cells in 2 of the 3 surviving TAM-treated *K5CreERT2^wt/mut^; Atg7^loxp/loxp^* mice that had been administered 4NQO (**Figure 9E**). By contrast, BCH was observed in only 2 of the 5 vehicle-treated *K5CreERT2^wt/mut^; Atg7^loxp/loxp^* mice that had been administered 4NQO (**Figure 9E**), none of which displayed a change in body weight (**Figure 9D**). As expected, no pathological alterations were identified in the absence of 4NQO treatment (**Figure 9D, E**). These findings suggest that autophagy inhibition in squamous epithelium *in vivo* is deleterious in the context of challenge with the carcinogen 4NQO.

**Figure 9.**
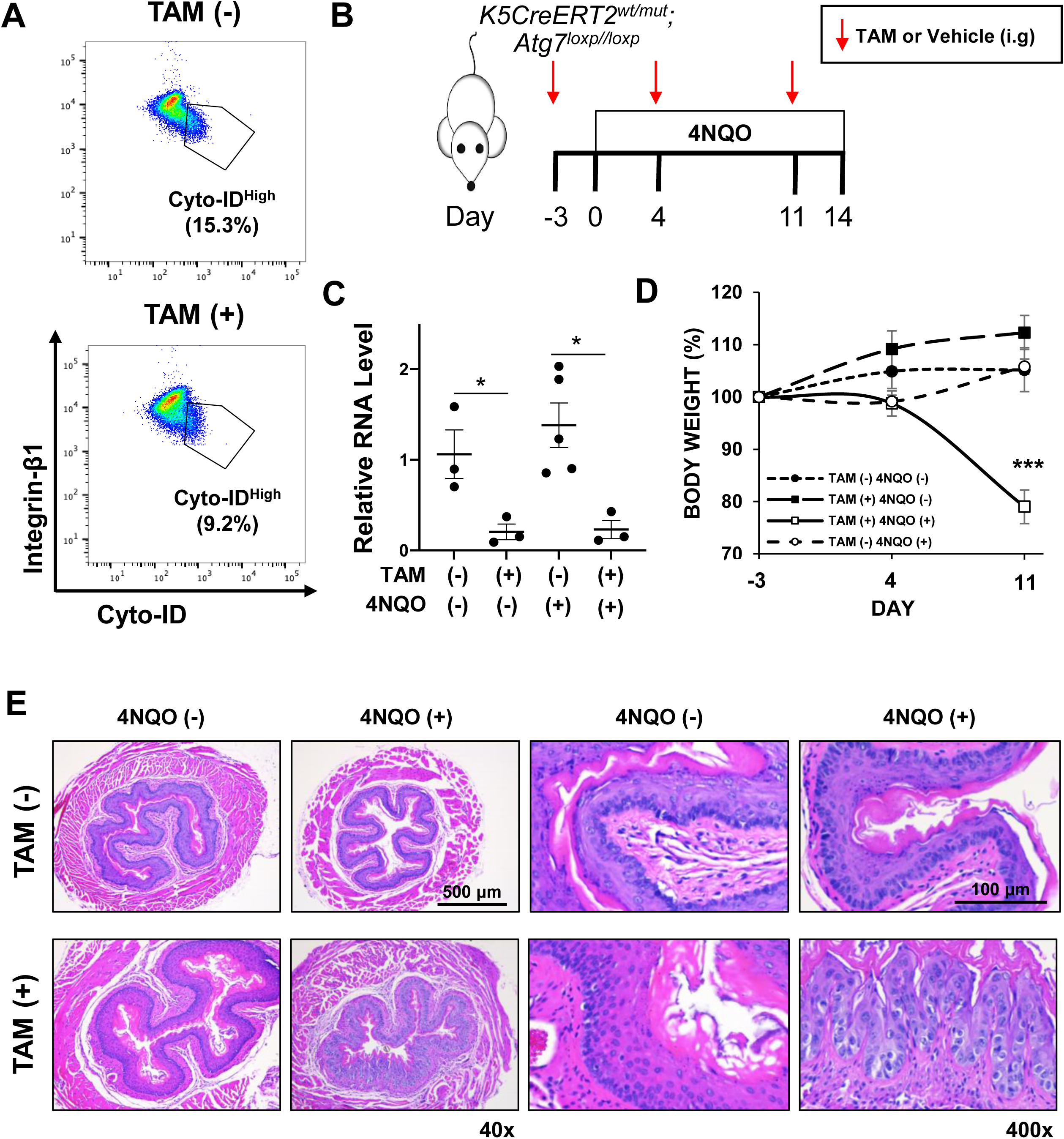
Effects of genetic Atg7 depletion in esophageal epithelium on response to carcinogen 4-nitroquinoline 1-oxide. (A) K5-Cre^ERT2+/-^ ; Atg7^loxp/loxp^ mice treated with tamoxifen (TAM) for 72 hours or vehicle-treated controls were evaluated for Cyto-ID^High^ cells in esophageal basal cells. **(B-E)** K5Cre^ERT2+/-^; Atg7^loxp/loxp^ mice were gavaged with vehicle or tamoxifen (TAM) at indicated time points. 4-nitroquinoline 1-oxide was administered for 14 days. Schematic overview of experimental design is shown in **(B).** Dot plot showing relative RNA expression of *Atg7* in epithelium-enriched portion of esophagus is shown in **(C)** with each dot representing an individual mouse and bars representing Mean±SEM (*p<0.05 by t-test by ANOVA). Body weight curves are shown in **(D)** (***p<0.0001). Representative H&E-stained sections from esophagi of mice in each experimental and control group are shown in (**E**).

## Discussion

Autophagy is a dynamic cellular process that has cell type- and context-dependent functions in diverse tissue types. In the esophagus, we and others have demonstrated roles for autophagy in pathological conditions both benign and malignant. Although such studies suggest the potential for broad functional significance of this evolutionarily conserved pathway in esophageal biology, what role—if any—autophagy plays in the esophagus under homeostatic conditions has remained elusive. Here, we address this critical knowledge gap, demonstrating that genetic depletion of the essential autophagy gene *Atg7* results in disruption of the defined esophageal epithelial proliferation-differentiation gradient, including increased basal cell proliferation. While genetic depletion of autophagy has been shown to limit proliferation of epidermal keratinocytes *in vitro,* the role of autophagy in squamous tissues *in vivo* is more complex. In the epidermis, conditional depletion of ATG7 in basal cells has been shown to increase thickness in the skin of the back with no effect on differentiation noted^30^. An independent study, however, identified epidermal differentiation defects in ATG7 depleted mice^31^. Conditional depletion of ATG7 or ATG5, an essential autophagy gene that facilitates AV biogenesis along with ATG7^32^, in basal cells has been shown to perturb differentiation in the preputial gland but have no effect on morphology of the thymus^33, 34^. In the skin and thymus, autophagy in basal cells has been defined as dispensable for organ function under homeostatic conditions *in vivo*. We failed to observe evidence of weight loss or food impaction in TAM-treated *K5CreERT2^wt/mut^; Atg7^loxp/loxp^* mice; however, direct measurement of esophageal barrier function and food transit is necessary to determine if ATG7-dependent autophagy is dispensable for esophageal function. Moreover, although Cyto-ID^High^ cells are reduced in TAM-treated *K5CreERT2^wt/mut^; Atg7^loxp/loxp^* mice, a model in which these cells are fully abrogated is necessary to define the dispensability of autophagy in esophageal epithelium.

Our *in vivo* studies further suggest that autophagy is critical for response to the oral- esophageal carcinogen 4NQO. We unexpectedly observed death and weight loss in mice with squamous epithelial-specific ATG7 depletion following 4NQO administration. Although 4NQO is a well-established carcinogen that promotes robust tumorigenesis in the oral cavity and esophagus with exposure over several months, acute effects of 4NQO are not well-characterized. Here, we find that 4NQO induces weight loss upon acute exposure and this is exacerbated with ATG7 depletion. Expression of autophagy markers, including ATG7, increases during 4NQO- mediated carcinogenesis in the oral cavity^35^; however, this has not been studied in the context of esophageal carcinogenesis. Following 4NQO exposure, esophageal epithelium of mice with squamous epithelial-specific ATG7 depletion displayed evidence of BCH and altered basal cell morphology that was defined as cytoplasmic vacuolization, a finding associated with pathogen-induced cell death in mammalian cells^36^. Xenophagy is a selective form of autophagy that contributes to host defense via targeting pathogens to AVs where they are degraded upon AV fusion with lysosomes.^37–39^. Future studies will determine if 4NQO administration in TAM-treated *K5CreERT2^wt/mut^; Atg7^loxp/loxp^* mice promotes pathogen accumulation in esophageal epithelium to drive weight loss and death. As activation of the K5 promoter is not confined to esophageal epithelium, it is important to consider that effects of autophagy depletion in other squamous tissues may contribute to the deleterious effects seen with 4NQO administration.

Our findings in *K5CreERT2^wt/mut^; Atg7^loxp/loxp^*mice along with our published work demonstrating that there is a gradient of staining for cleaved LC3, a marker of AVs, in murine esophageal basal cells^23, 24^, led us to investigate whether intracellular AV content is associated with molecular and functional attributes of esophageal basal cell. Experiments in 3D organoids generated by sorted Cyto-ID^Low^ and Cyto-ID^High^ cells supported this premise with Cyto-ID^High^- derived organoids being less capable of generating 3D organoids upon initial plating but then displaying increased organoid passaging capability indicative of self-renewal. Coupled with self-renewal, additionally, we found the transcriptional profiles of Cyto-ID^Low^ and Cyto-ID^High^ cells to be markedly different with more than 7,000 DEGs. To put this finding into context, our previous study comparing the transcriptional profiles of *Krt15*-positive and *Krt15*-negative esophageal basal cells revealed <200 DEGs^16^. Consistent with impaired ability to generate organoids, Cyto-ID^High^ cells that were freshly isolated from murine esophageal epithelium exhibited diminished proliferation and compared to their Cyto-ID^Low^ counterparts. By contrast, organoids generated by Cyto-ID^High^ cells displayed increased cell proliferation as compared to Cyto-ID^Low^-derived organoids. As autophagy is often activated as a stress response, it is not surprising that cell cycle arrest has been shown to positively correlate with autophagy in mammalian cells^19^. Investigation of the mechanisms that drive the switch in Cyto-ID^High^ cells from a low proliferative state in the esophagus to a higher proliferative state in 3D organoids will be of great interest. As 3D organoids generated Cyto-ID^High^ cells have similar AV content to those generated by Cyto-ID^Low^ cells, autophagy may not be required for increased proliferation and self-renewal in Cyto-ID^High^-derived organoids. This must be formally tested, however. Autophagy has been implicated in paligenosis, a process through which fully differentiated cells re-enter the cell cycle^40, 41^. Given that organoids generated Cyto-ID^High^ cells display aberrant proliferation in cells located beyond the basal cell compartment, examination of paligenosis in these organoids is warranted. Paligenosis occurs in response to cell stress which may indeed be induced as esophageal epithelial cells are cultured in *ex vivo* 3D organoid culture.

The molecular and functional attributes of Cyto-ID^High^ esophageal basal cells are consistent with a basal population that exhibits limited proliferation coupled with stem-like properties, namely the ability to self-renew and give rise to terminally differentiated progeny. There has been great interest—and controversy—in examining basal cells as esophageal stem/progenitor cells. Pan et al. tracked 5-iodo-2’-deoxyuridine label-retaining cells with features of stem cells (long-lived, slow-cycling, uncommitted, multipotent) residing primarily in the basal cell layer of normal human esophageal epithelium^42^. Consistent with the presence of a slow-cycling label-retaining cells with self-renewal capacity have been identified in the esophageal basal layer where they are marked by positivity for CD34 or a combination of high expression of integrin α6 and low expression of CD71^5, 17^. Giroux et al. further demonstrated that long-lived K15- positive basal cells display self-renewal and tissue regeneration capacity^16^. We have recently identified a population of slow-cycling, CD73-positive cells in the basal and suprabasal layers of human esophageal epithelium that contributes to epithelial renewal^43^. By contrast, work by DeWard et al. suggested that an actively proliferating subset of basal cells defined by positivity for CD73 and high expression of integrin β4 exhibits the greatest stem cell potential in murine esophageal epithelium^18^. In light of the described studies, it is tempting to speculate that basal cells with high AV content represent a novel slow-cycling stem/progenitor population in esophageal epithelium. Such a conclusion, however, is at odds with studies utilizing lineage tracing in mice coupled with mathematical modeling to propose a single-progenitor model wherein all basal cells have equal capacity to proliferate and differentiate^1, 19^.

As autophagy is a dynamic cellular process it quite possible that AV level is reflective of a cell state rather than serving as a marker for a defined subset of esophageal basal cells. We propose a model (**Figure 10**) wherein esophageal basal cells have high levels of AVs when they are directed by internal and/or external cues to suppress proliferation. These same cells may then decrease their AV context should the need to proliferate and/or differentiate arise. When we take Cyto-ID^Low^ cells from their local tissue microenvironment, they are poised to proliferate and differentiate, thus giving rise to typical esophageal organoids. By contrast, Cyto-ID^High^ cells are in a state of cell cycle arrest when they are exposed to growth factor-rich 3D organoid media. In this context, Cyto-ID^High^ cells re-enter the cell cycle, albeit in an aberrant fashion with cells outside of the basal cell compartment exhibiting markers of proliferation, and also acquire self-renewal capability. In relation to our *in vivo* studies in *K5CreERT2^wt/mut^; Atg7^loxp/loxp^* mice, depletion of ATG7 diminishes Cyto-ID^High^ cells, allowing expansion of proliferative Cyto-ID^Low^ cells and this phenotype is likely exacerbated with 4NQO challenge. It is further possible that 4NQO may promote a local microenvironment that activates the self-renewal potential of Cyto-ID^High^ cells, similar to our findings in 3D organoids. In the intestine, we have recently demonstrated that high AV level prospectively identifies facultative stem cells that contribute to the DNA damage response following ionizing radiation exposure^44^. It remains to be determined if facultative stem cells marked by high AV level contribute to esophageal response to stressors including 4NQO.

**Figure 10.**
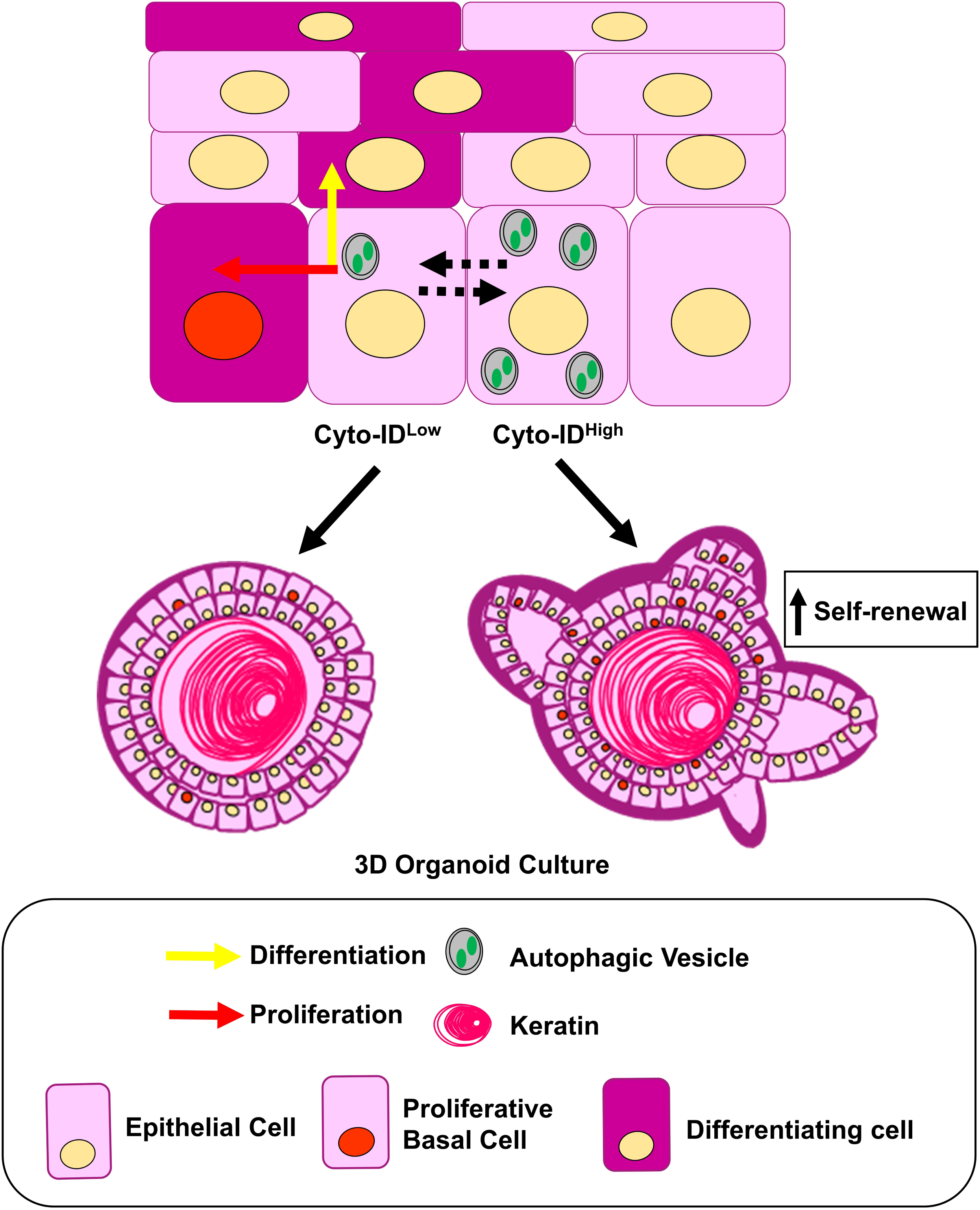
Proposed model. In murine esophageal epithelium, basal cells with high AV levels (Cyto-ID^High^) are largely non-proliferative while those with low AV levels (Cyto-ID^Low^) readily proliferate, stochastically giving rise to basal cells or executing squamous differentiation. Basal cells with high and low levels of AVs may interconvert. When we take Cyto-ID^Low^ cells from their local tissue microenvironment, they are poised to proliferate and differentiate, thus giving rise to typical esophageal organoids. When Cyto-ID^Low^ are placed into 3D organoid culture they continue to proliferate and differentiate. When growth-arrested Cyto-ID^High^ cells are placed into 3D organoid culture they exhibit aberrant proliferation yet remain capable of differentiating. Cyto-ID^High^ cells also gain self-renewal capability in 3D organoid culture.

In summary we report previously unappreciated roles for autophagy in esophageal homeostasis and response to 4NQO challenge. We show that esophageal basal cells with high AV level exhibit limited proliferation yet have the ability to self-renewal in the context of *ex vivo* 3D organoid culture. As these studies are solely conducted in mice, it will be of great interest to determine the relevance of Cyto-ID^High^ esophageal basal cells in human physiology. Notably, we have previously demonstrated that esophageal epithelial cells with Cyto-ID fluorescence can be detected in human subjects with normal esophageal pathology and that Cyto-ID fluorescence increase in EoE patients^23^. Our understanding of the mechanisms regulating esophageal basal cell remains quite limited despite the potential of this knowledge to guide novel approaches for diagnosis, monitoring, and therapy of widely prevalent esophageal diseases. Autophagy is a particularly attractive candidate in this regard given that several FDA-approved drugs have been shown to impact this pathway.

## Materials and Methods

### Murine Studies

All murine studies were performed in accordance with a protocol approved by the Temple University IACUC (Protocol Number: 5018). Experiments were conducted in accordance with institutional guidelines for animal research. Atg7^loxp/loxp^ (Strain# RBRC02759; RIKEN BRC), *K5CreERT2^wt/mut^* (Strain# 029155; Jackson Labs) and C57BL/6 (Strain# 007909; Jackson Labs) mice were maintained under controlled conditions with a 12 h light/dark cycle. *K5CreERT2^wt/mut^; Atg7^loxp/loxp^* mice were administered TAM (250 μg/g body weight dissolved in peanut oil) by oral gavage to induce Cre-mediated excision of exon 14 of ATG7 in cells in which the Krt5 promoter is active^45^. Vehicle controls, designated TAM (-), were gavaged with an equal volume of peanut oil. 4NQO (100 μg/ml in 2% propylene glycol was administered via drinking water. Vehicle controls, designated 4NQO (-), were administered 2% propylene glycol. Body weight of mice was monitored at least once/week during all experiments. All treatments were initiated in 6-20-week-old mice. Number of animals in each experimental group are noted in corresponding figure legends. Both males and females were included in all animal studies.

### Flow Cytometry & Cell Sorting

Murine esophagi were dissected from mice and peeled epithelial-enriched layer was enzymatically digested^11, 12^. To isolate Cyto-ID^Low^ and Cyto-ID^High^ esophageal basal cells, the resulting single cell suspension was incubated at 37°C for 30 minutes with Integrin β1 antibody (APC anti-mouse/rat CD29; 102216; Biolegend) at 1:100 dilution. Cells were washed, pelleted at 1000 RPM for 5 minutes, and incubated at 37°C for 30 minutes with Cyto-ID (ENZ-51031-K200; Enzo Life Sciences) at 1:1000 dilution. After washing, cells were resuspended in FACS buffer (1% of BSA in PBS) and 2 μg/ml DAPI (D1306; Invitrogen) was added immediately before sorting using FACS Aria Cell Sorter (BD Biosciences). Total live esophageal epithelium cell compartment (DAPI^negative^; Integrin β1^positive^) was sorted based upon Cyto-ID fluorescence into two populations: Cyto-ID^Low^ and Cyto-ID^High^. For image flow cytometry, single cell suspension from mouse esophagi was incubated at 37°C for 30 minutes with Integrin β1 antibody (PE anti-mouse/rat CD29; 102207; Biolegend) at 1:200 dilution. Then cells were stained with Cyto-ID as described above. 2 μg/ml DAPI was added immediately before analysis by Amnis ImageStream imaging flow cytometer, Program Ideas version 6.2. Analysis was performed by trained staff at the University of Pennsylvania Flow Cytometry Core. For cell cycle analysis, single cell suspension from mouse esophagi or 3D organoids fixed for at least 2 hours at 4°C using 66% ice cold ethanol in PBS. After fixation, cells were equilibrated to room temperature and stained with PI for 30 minutes at 37°C using Propidium Iodide Flow Cytometry Kit (ab139418; Abcam) according to the manufacturer’s protocol and incubated at 37°C in the dark for 30 minutes. Flow cytometric analysis was done using a BD LSR II (BD Biosciences) and analyzed with FlowJo software (Tree Star).

### Electron Microscopy

Electron microscopy was performed as previously described^23, 24, 46^. Briefly, sorted Cyto-ID^Low^ and Cyto-ID^High^ cells were fixed in cacodylate-buffered 2.5% (w/v) glutaraldehyde, post-fixed in 2.0% osmium tetroxide, and then embedded in epoxy resin and ultrathin sections post-stained in the University of Pennsylvania Electron Microscopy Resource Laboratory. Images were obtained using a JEOL-1010 transmission electron microscope fitted with a Hamamatsu digital camera and AMT Advantage imaging software.

### 3D Esophageal Organoid Assays

Sorted Cyto-ID^Low^ and Cyto-ID^High^ murine esophageal basal cells were resuspended in mouse KSFM medium and mixed with 90-95% Matrigel (354234; Corning). Using 24 well plates, 500- 1000 cells were seeded per well in 50 μl Matrigel. After solidification, 500 μl of Advanced DMEM/F12 (12634-101; Gibco) supplemented with 1X Glutamax (35050-061; Gibco), 1X HEPES (15630-080; Gibco), 1% v/v penicillin-streptomycin (15140-122; Gibco), 1X N2 Supplement (17502-001; Gibco), 1X B27 Supplement (17504-044; Gibco), 0.1 mM N-acetyl-L-cysteine (616-91-1; Fisher Chemical), 50 ng/ml human recombinant EGF (10450-013; Gibco), 3.0% Noggin/R-Spondin-conditioned media, and 10 μM Y27632 (1254; Tocris Bioscience) was added to each well. After 13-15 days of culture, organoids were imaged and collected for histology and/or passaging. For passaging, single cells were isolated from organoids via incubation for 5 minutes with 1X Dispase (354235; Corning) in HBSS followed by incubation for 1 hour at 37°C with shaking at 1000 RPM in 0.25% Trypsin-EDTA (25-510; Genesee Scientific) supplemented with 10 μM Y27632. Cells were then forced through a cell strainer (70 μm) into a tube containing 4 ml 250 μg/ml soybean trypsin inhibitor (17975-209; Gibco) in PBS. Cells were pelleted then resuspended in complete ‘mouse’ keratinocyte serum-free medium (KSFM): KSFM without calcium chloride (10725-018, Gibco, Waltham, MA, USA) with recombinant epidermal growth factor (1 ng/ml), bovine pituitary extract, (50 mg/ml), 1% penicillin/streptomycin (15140-122; Gibco, and 0.018 mM CaCl_2_ (349610025; Acros Organics) . Single cells were then replated into Matrigel. For single cell RNA-Seq of dissociated 3D organoids, single cell suspension was enriched for live cells using Dead Cell Removal kit (130-090-101; Miltenyi Biotec) according to the manufacturer’s protocol.

### Histological Analysis

Whole esophagi were dissected and fixed in 10% neutral buffered formalin for 12 hours at 4°C. Tissues were washed with PBS then stored in 70% ethanol at 4°C prior to paraffin embedding. Organoids were grown for 12-15 days before recovering from Matrigel with Dispase I and fixing overnight in 4% paraformaldehyde (J19943-K2; Thermo Scientific). Specimens were embedded in 2.0% Bacto-Agar: 2.5% gelatin prior to paraffin embedding. Sectioning and hematoxylin and eosin (H&E) staining were performed at the Fox Chase Cancer Center Histopathology Core. Immunohistochemical staining was performed on a VENTANA Discovery XT automated staining instrument (Ventana Medical Systems) using VENTANA reagents according to the manufacturer’s instructions. Tissue sections were incubated with anti-ATG7 antibody or anti-Ki67 antibody overnight at 4°C followed by incubation with corresponding secondary antibody. No primary antibody was utilized as a negative control for ATG7 staining and normal rabbit IgG was used as a negative control for Ki-67 staining. Samples were developed using DAB Substrate Kit,

Peroxidase (with Nickel) (SK-4100, Vector Laboratories) or VENTANA ChromMap DAB detection kit (760-159). Immunostained slides were scanned using an Aperio ScanScope CS 5 slide scanner (Aperio, Vista, CA, USA) and were viewed with Aperio’s image viewer software (ImageScope, version 11.1.2.760, Aperio). Selected regions of interest were outlined manually by a pathologist (AKS, KQC). The Ki-67 quantifications were performed with Aperio V9. 4 LOIs (lesions of interest) were selected for each case. Pathologist (AKS, KQC) blinded to treatment parameters quantified Ki67 staining as percentage of Ki67-positive basal cells out of 200-300 and measured thickness of esophageal epithelium. Slides were imaged using Leica DM 1000 LED microscope. A list of antibodies with used dilutions is provided in **Table 1**.

### Quantitative Real-Time PCR

Epithelium-enriched portion of murine esophagus was transferred to 1 ml RNAlater Solution (AM7020; Invitrogen) and stored at room temperature for up to 24 hours. Tissue was homogenized on ice in RLT buffer (74104; Qiagen, Hilden, Germany) with 1% β-mercaptoethanol (444203; EMD Millipore Corporation) and RNA was isolated by using RNeasy Mini Kit (74104; Qiagen) according to the manufacturer’s protocol. cDNA was synthesized from approximately 100-300 ng total RNA using the Maxima First Strand cDNA Synthesis Kit for RT-qPCR (K1642; Thermo Fisher Scientific) according to the manufacturer’s protocol. qPCR performed was using SYBR Green Master Mix (A46109; Applied Biosystems) and was analyzed using MiniAmp Plus Thermal Cycler (A37835; Applied Biosystems). Gene targets and primer sequences used are listed in **Table 2**.

### Bulk RNA-Seq & Data Analysis

RNA was isolated from sorted Cyto-ID^Low^ and Cyto-ID^High^ murine esophageal basal cells using RNeasy Mini Kit (74104; Qiagen) according to the manufacturer’s protocol. Library construction, quality check, quantification, and RNA-Seq were performed by Genomics Facility at Fox Chase Cancer Center. Briefly, to make stranded mRNA-Seq library: 500 ng total RNA from each sample were used to make library according to the product guide of stranded mRNA library kit (E4720L; NEBNext Ultra™ Directional RNA Library Prep Kit for Illumina). In short, mRNAs were enriched twice via poly-T based RNA purification beads and subjected to fragmentation at 94^°^C for 15 minutes via divalent cation method. The 1^st^ strand cDNA was synthesized by reverse transcriptase and random primers at 42^°^C for 15 minutes, followed by 2^nd^ strand synthesis at 16^°^C for 1 hour. A single ‘A’ nucleotide was added to the 3’ ends of the blunt fragments at 37^°^C for 30 minutes. Adapters with IlluminaP5, P7 sequences as well as indices were ligated to the cDNA fragment at 30^°^C for 10 minutes. After purification using SPRIselect Beads (B23318;Beckman Coulter), a 13-cycle PCR reaction was used to enrich fragments. PCR was set at 98^°^C for 10 seconds, 60^°^C for 30 seconds, and extended at 72^°^C for 30 seconds. Libraries were again purified using SPRIselect Beads. Quality check was performed on Agilent 2100 bioanalyzer using Agilent high sensitive DNA kit (5067-4626). Quantification was performed with Qubit 3.0 fluorometer (Q33216, ThermoFisher Scientific) using Qubit 1X dsDNA HS Assay Kit (Q33230). Sample libraries were subsequently pooled and loaded to Nextseq 500. Samples were sequenced 50-200 bases pair-end on Nextseq 500 machine with 30 million read depth. FASTQ files were obtained at Illumina Base Space. Sequencing reads were aligned to the mouse genome GRCm38 and gene counts were calculated using STAR v2.7.3 aligner. Differential expression analysis was performed in R using the DESeq2 package. DESeq2 calculates differential expression by median of ratios method. DESeq2 calculated fold changes and significance. Sample-specific gene ratios are calculated as the gene count over the geometric mean for each specific gene. P-values are adjusted for multiple testing by Benjamin-Hochberg. Then, Principal Component Analysis (PCA) was used to visualize the samples’ grouping according to their sex and Cyto-ID status. After alignment and normalization (calculating relative values based on the total amount of counts in the sample), we found 55,276 genes, among which 8,641 genes were significantly differentially expressed in Cyto-ID^High^ cells compared to Cyto-ID^Low^ cells (padj-value <0.05). To improve the quality of data processing we further analyzed genes with padj <0.01. This resulted in 7,102 genes. These genes were visualized by Heatmap using Pheatmap package and pathway analysis was performed using Ingenuity Pathway Analysis (IPA; Qiagen) core and comparison analysis. Without adding additional cutoffs, 6,782 of 7,102 were mapped and resulted in 110 significantly dysregulated pathways. The top 10 most significantly up and downregulated IPA pathways were ranked based on z-score.

### scRNA-Seq & Data Analysis

3D organoids were dissociated to generate single cell suspension as described above. Single cell droplets were generated with chromium single-cell controller using Chromium Next GEM Single Cell 3’ Kit v3.1 (1000121; 10x Genomics). 5000–7000 cells were collected to make cDNA at the single cell level. Full-length cDNA with unique molecular identifiers (UMIs) was synthesized via reverse-transcription in the droplet. After PCR amplification and purification, cDNA was fragmented to around 270 bp and Illumina adapters with index were ligated to fragmented cDNA. After PCR, purification, and size selection, single cell RNA libraries were 450 bp in length and sequenced on Illumina sequencer at R1 = 28 bp, R2 = 91 bp.

Deconvolution of scRNA-Seq reads followed 10X Genomics Cell Ranger (v6.0.0) pipeline^47^. Massively parallel digital transcriptional profiling of single cells was performed using command ‘cellranger count’ with FASTQ files as input from each sample. For cellranger count, R1 and R2 were trimmed to 28 and 91 bp, respectively, to remove PCR adapters. The mouse genome mm10 (GENCODE vM23/Ensembl 98) was used as the reference for genome alignment and feature counting. From the output, the filtered matrices were used for downstream analyses. Matrices for each murine peeled epithelium sample were imported and transformed into Seurat (v4) objects for further processing. Cells with over 10% transcripts consisting of mitochondrial genes, under 1,500 unique genes, over 6,000 unique genes, or over 60,000 total UMIs were excluded to remove doublets or dead/dying cells. 13,667 cells passed this threshold. Analysis of the filtered matrices follows Seurat integration workflow described by Stuart, et. al.^48^ using SCTransform function, which normalizes counts while accounting for read depth and subsequently searches for the top 2,000 most variable feature per sample with corrected counts for integration. Reciprocal PCA was then used to find integration transcript anchors between all matrices. Genes used for integration were ranked by the number of matrices they appear in. From this point on, dimensionality reduction used genes and values that were pre-processed using the integration workflow. However, raw and normalized counts were stored for downstream differential expression tests. The resulting dataset was then reduced via PCA, resulting in 30 principal components. An elbow plot was generated to examine the standard deviations of each component, which verified that the first 30 principal components contain most of the sources of variation in the dataset. To capture all the sources of variation in the dataset, all principal components were then used as input to the UMAP dimensionality reduction procedure (arXiv:1802.03426). Because of our interest in the relationship between cell cycle phases and cell fates, we opted not to regress cell cycle genes in our dimensionality reduction steps. A Shared Nearest Neighbor (SNN) graph was then constructed with the principal components of PCA by first determining the nearest neighbors for each cell and subsequently creating the SNN guided by neighborhood overlap between each cell and its nearest neighbors. Clusters were then determined by a modularity optimization algorithm by Waltman and van Eck^49^. To choose an optimal clustering resolution, a clustering tree was generated with the R package cluster which depicts movement of cells across clusters as resolution is increased from 0.1 to 1 with 0.1 increments. Resolution 0.4 was chosen, as it is the earliest resolution that created several clusters that were stable as the resolution was increased, as well as having a minimal number of clusters that were composed of multiple clusters from the next lowest resolution.

For each cluster, differentially expressed genes (DEGs) were calculated by comparing expression of genes within the cells of the cluster over the expression of the genes in all other clusters. The statistical workflow to determine differential expression was Seurat’s implementation of Wilcoxon’s signed-rank test. Significance cutoff for DEGs is a Bonferroni-adjusted p value of 0.05, and the fold change cutoff is below −0.25 or above 0.25 natural log fold change. To compare each clusters’ proportional size between the two Cyto-ID groups, each sample’s cluster proportions were calculated, and a Wilcoxon signed-rank test was performed to compare mean cluster proportions between the two Cyto-ID groups. Log normalized expression of *Krt5* vs. normalized log_10_ expression of *Krt13* expression was plotted for all populations identified in the single cell RNA-Seq dataset. Populations falling within the strip defined by lines with a slope of 1 passing through points (0, −15) and (0, 15) were defined as suprabasal. Populations falling below this strip were defined as basal and those above this strip were identified as superficial. Cell populations were imported into QIAGEN IPA for core analysis.

The Seurat function CellCycleScoring was used to predict cell cycle phase of each cell. The function takes as input a list of S phase upregulated genes and G2/M upregulated genes, and outputs the score for each phase. The S and G2/M phase genes were provided within the Seurat package as objects “cc.genes.updated.2019$s.genes” and “cc.genes.updated.2019$g2m.genes”, respectively. The cell cycle phase is determined by the dominating score. Cells with weak scores for both phases are classified as G0/G1 phase cells. To compare each clusters’ proportional size between the different Cyto-ID groups, each sample’s cluster proportions were calculated, and a Wilcoxon signed-rank test was performed to compare the mean cluster proportions between the two Cyto-ID groups.

### Statistical Analysis

Statistical analyses were performed as described in corresponding figures. Tests were performed using R and GraphPad Prism (GraphPad Software, La Jolla, CA, USA). Analysis of variance (ANOVA) was performed on all experiments with more than 2 groups. p<0.05 was used as the threshold for statistical significance. All experiments presented were conducted at least in triplicate to ensure reproducibility of results.

## Supporting information

Supplementary Table IPA Fig 3c

Supplementary Table IPA Fig 7

Supplementary Table IPA Fig 3b

Supplementary Table IPA Fig 5c

## Acknowledgements

We thank David Ambrose and Amir Yarmahoodi for assistance with flow cytometry and cell sorting at the Lewis Katz School of Medicine Flow Cytometry Core. We thank staff at the University of Pennsylvania Electron Microscopy Resource Laboratory and University of Pennsylvania Flow Cytometry Core for technical assistance. This work was supported by the following grants: R01DK121159 (KAW), R01DK121159-S1 (JLJ), T32GM142606 (ADF; PIs: Xavier Graña, Jonathan Soboloff, Temple University), P30CA006927 (YT, KCC, AKS; PI: Jonathan Chernoff, Fox Chase Cancer Center).

## Data availability statement

All sequencing data and associated metadata has been deposited into GEO. At the time of publication, all data generated in this study will be available from online repositories with accession IDs GSE243539 and GSE243580 or can be obtained upon request from the corresponding author.

## Author Contributions

AK: project administration, contribution to the design of the study, data curation, formal analysis and writing—original draft; ALK: statistical analysis; ADF: statistical analysis, writing—review and editing; LRP: collection of the data; SRP: collection of the data; SN: collection of the data; JLJ: writing—review and editing; AM: collection of the data; YT: collection of the data; KQC: formal analysis; AKS: formal analysis; ABM: conceptualization, supervision and writing—reviewing and editing; MPT: conceptualization, supervision and writing—reviewing and editing; KEH: conceptualization, supervision and writing—reviewing and editing; KAW: conceptualization, initiation of the study, supervision and writing—reviewing and editing. All authors contributed to the article and approved the submitted version.

