## Supplementary Figure 1 for "Autophagy contributes to homeostasis in esophageal epithelium where high autophagic vesicle content marks basal cells with limited proliferation and enhanced self-renewal potential"

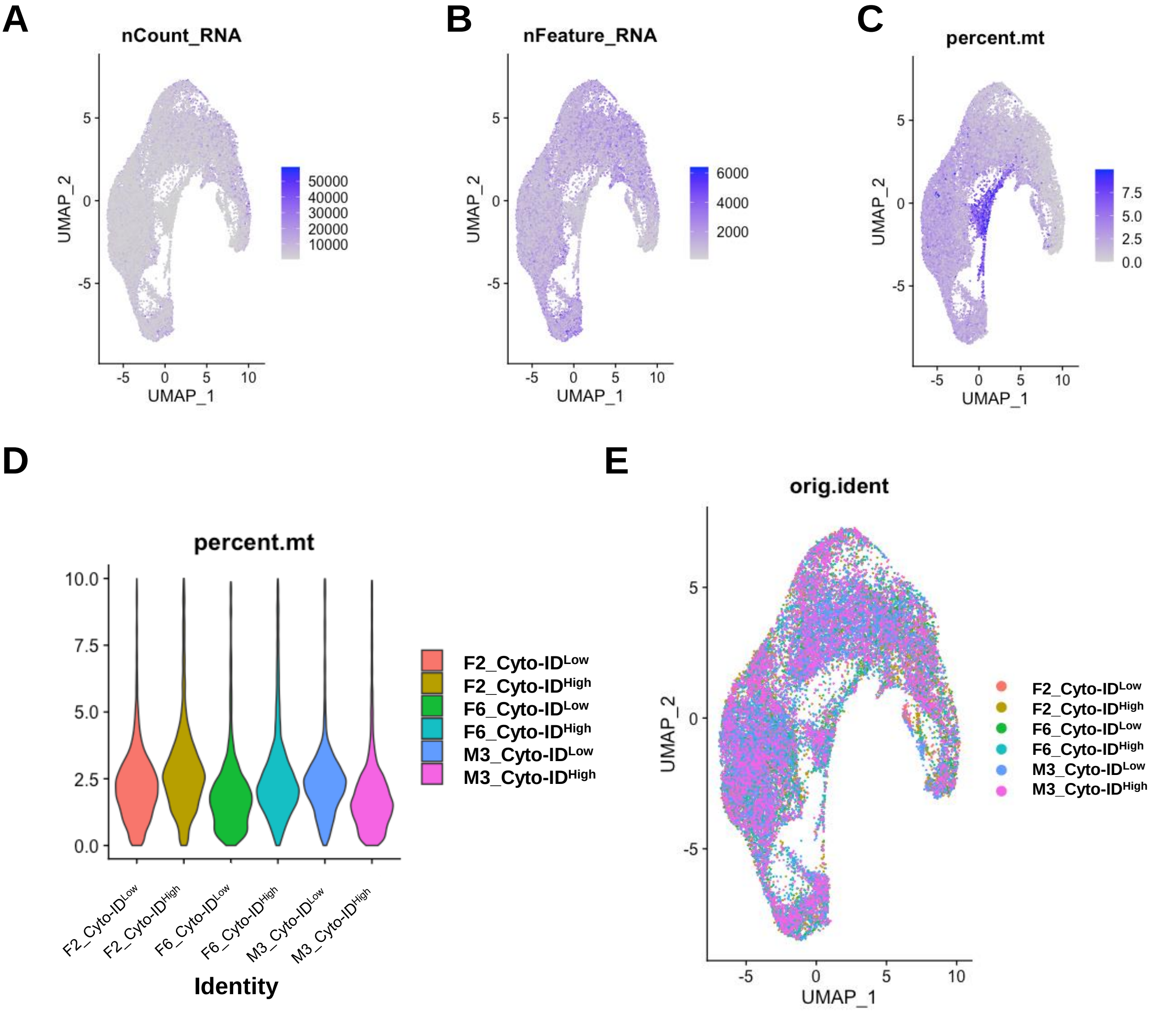


**Supplementary Figure S1. Quality control metrics for single-cell RNA sequencing data. (A)** Transcript counts for all samples and distribution across the uniform manifold approximation and projection (UMAP) object are shown. **(B)** Unique transcript counts for all samples and distribution across the UMAP object is shown. **(C)** Percent expression of mitochondrial genes for all samples and distribution across the UMAP object is shown. **(D)** Violin plots with sample-based mitochondrial genes expression distribution. **(E)** Each cell is colored according to sample from which from it was derived and distribution across the UMAP object is shown.
